# Training Genotype Callers with Neural Networks

**DOI:** 10.1101/097469

**Authors:** Remi Torracinta, Fabien Campagne

## Abstract

We present an open source software toolkit for training deep learning models to call genotypes in high-throughput sequencing data. The software supports SAM, BAM, CRAM and Goby alignments and the training of models for a variety of experimental assays and analysis protocols. We evaluate this software in the Illumina Platinum whole genome datasets and find that a deep learning model trained on 80% of the genome achieves a 0.986% accuracy on variants (genotype concordance) when trained with 10% of the data from a genome. The software is distributed at https://github.com/CampagneLaboratory/variationanalysis. The software makes it possible to train genotype calling models on consumer hardware with CPUs or GPU(s). It will enable individual investigators and small laboratories to train and evaluate their own models and to make open source contributions. We welcome contributions to extend this early prototype or evaluate its performance on other gold standard datasets.

## INTRODUCTION

We recently presented an approach to call somatic variations with deep learning models, using no gold standard variations (Torracinta et al. [2016], Campagne [2016]). In this study, we use high-throughput sequencing data and available labels from gold-standard datasets to train deep-learning models that can call genotypes. We built on the variationanalysis software we presented in Torracinta et al. [2016], Campagne [2016] and distribute release 1.2 of the project, which now also supports training and using genotyping models. In this preprint, we present the methods we used to train and evaluate genotyping models, discuss how our approaches differ from the recently presented study of Pollin et al, and present evaluation data on the Illumina platinum genome data.

The methods we present rely on deep learning models that are trained from data. These methods are general and can be adapted to the characteristics of new experimental or analysis protocol by retraining models with sequence data obtained with commercially available DNA samples Torracinta et al. [2016], Campagne [2016], Poplin et al. [2016]. We present a brief protocol demonstrating how deep learning models can be trained with the open-source variationanalysis project. To our knowledge, our study is the first to offer an open-source implementation of a deep learning genotype caller and provide automated protocols to train models for new assays. The methods developed for this caller support diploid as well as polyploid organisms.

## RESULTS

We developed a deep learning genotype caller. Briefly, the caller uses the Goby framework (Campagne et al. [2013]) to observe characteristics of read alignments against a reference genome, and the variationanalysis project (Torracinta et al. [2016], Campagne [2016]) to vectorize these characteristics into features and labels suitable for training a feed forward neural network.

The caller can be trained using alignments in BAM, CRAM or Goby formats (Li et al. [2009], Fritz et al. [2011], Campagne et al. [2013]) and associated labels. Labels necessary to train neural networks are obtained from gold-standard datasets. Here, we used the Illumina Platinum Genomes (Eberle et al. [2016]) as a source of alignments and true genotypes. In this first study, we called SNPs and disregarded indels. All performance metrics are presented for SNPs.

We assembled a training dataset using all variants matching sites in the Platinum Genome NA12877 sample and 10% of other non-variant sites. This dataset was split in a training set (80% of sites), validation set (10% of sites, further sub-sampled on non-variant sites) and test set (10% of sites). The variationanalysis project provides tools to simplify assembling datasets (and make their production consistent). See Material and Methods for a summary of the protocol and project documentation online for details).

In this study, we mapped alignment data to 642 features for each site. Table 1 summarizes the characteristics of these datasets.

**Table 1.**
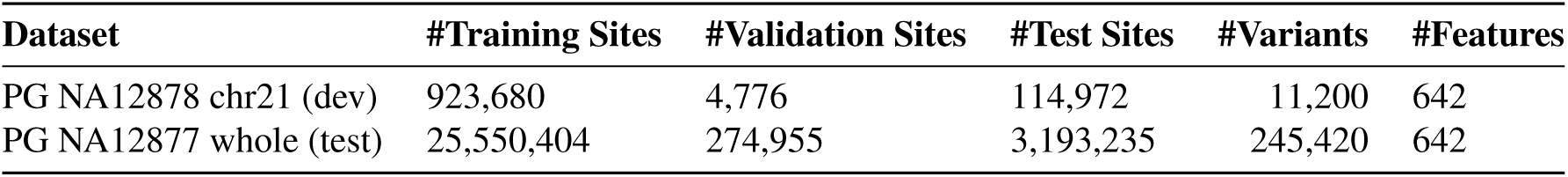
Characteristics of Datasets

| Dataset | #Training Sites | #Validation Sites | #Test Sites | #Variants | #Features |
| --- | --- | --- | --- | --- | --- |
| PG NA12878 chr21 (dev) | 923,680 | 4,776 | 114,972 | 11,200 | 642 |
| PG NA12877 whole (test) | 25,550,404 | 274,955 | 3,193,235 | 245,420 | 642 |

Initial feature mapper development was conducted on a development dataset composed exclusively of chromosome 21 alignments from NA12878. This smaller dataset is independent of the final training dataset since it was sequenced from a different individual. The purpose of feature mapper development is to identify a mapping from alignment data to feature and label vectors that result in predictive models on independent test datasets. We iteratively developed and tested about 15 mappers, identifying and fixing software bugs through error analysis after each iteration. Error analysis consists in examining the types of errors that the model makes (e.g., using a genome browser to visualize sites of prediction and the alignment) on the test dataset. This process often suggests features that should be presented as input to the network to facilitate learning. We stopped this process when performance seemed to reach a plateau on the small development set, suggesting that we needed more data to train the model.

Table 2 shows the performance metrics obtained on the development set with a reasonably tuned mapper. While these performance metrics are still far from the state of the art, they indicate that the mappers do a reasonably good job of mapping alignments to vectors since a reliable model can be trained with just one chromosome worth of data.

**Table 2.** Performance on Development Set. GC: Genotype Concordance. AUC: Area Under the Received Operating Curve for correct variant identification. Precision and Recall: estimated over variants only. F1: harmonic mean of Precision and Recall. Accuracy: estimated overall all genotypes.

| Dataset | Accuracy | Recall | Precision | F1 | GC | AUC |
| --- | --- | --- | --- | --- | --- | --- |
| chr21 dev | 0.997 | 0.977 | 0.924 | 0.950 | 0.977 | 0.889 |

The performance of deep neural networks are known to improve markedly when models are trained with larger numbers of training examples. To determine the improvement that more training data would bring, we trained a model with 10% of the data from the genome. Training was stopped when performance measured on the validation set did not increase after 10 epochs (complete passes over the training set). Table 3 shows the performance obtained when the model is trained with data from 10% of the genome (excluding 90% of non-variants containing sites, but keeping all variant containing sites). Performance metrics include all the gold-standard variants reported by the Platinum genome project (we did not limit this analysis to variant sites that overlap confident regions, but used all variants).

**Table 3.** NA12877: Performance on the Test Set. (10% of sites in the genome. The number of variants is the actual number of variants in the test set, used to estimate Precision, Recall and AUC. The AUC point estimate and 95% confidence intervals are estimated on a random sample of 50,000 of these sites.

| Accuracy | Recall | Precision | F1 | numVariants | GC | AUC | [AUC95 | AUC95] |
| --- | --- | --- | --- | --- | --- | --- | --- | --- |
| 0.994 | 0.986 | 0.936 | 0.961 | 225,941 | 0.986 | 0.908 | 0.904 | 0.911 |

These results show that more data can indeed markedly improve the performance of the trained model. These results place the open-source caller that we present within less than 1% of genotype concordance performance reported by Poplin and colleagues, while their models were training with data from complete genomes. Importantly, the recall of the models is close to optimal (0.986), while precision is still sub-optimal (0.936), suggesting that error analysis on the 10% dataset and future feature mapper optimizations could quickly bring performance in a competitive range.

## MATERIALS AND METHODS

### Alignment Data

Alignments were not pre-processed and were used directly after download from ftp://ussd-ftp.illumina.com/2016-1.0/hg19/small_variants/. Limited preprocessing was performed with Goby to realign variations around indels to eliminate mis-alignment artifacts. Minimal pre-processing is in contrast to Poplin et al. [2016] which used a haplotype realigner and several preprocessing steps that already cleaned up the data before providing it to the model.

### Source of Gold Standard Genotypes

We used all genotypes contained in the VCF files distributed at ftp://ussd-ftp.illumina.com/2016-1.0/hg19/small_variants/. In contrast to the study of Poplin et al. [2016], we did not restrict the gold standard variations to regions of high-confidence because this only trains the models on simpler regions of the genome and may limit its ability to discriminate variants in other regions.

### Software implementation

#### Source Code

The source code of the software used for this study is distributed under the open-source Apache 2.0 license at https://github.com/CampagneLaboratory/variationanalysis.

#### Using trained models

Models trained in this study are being integrated into release 3.2+ of the Goby genotype caller (distributed at http://github.com/CampagneLaboratory/goby3). Goby3 supports alignments in the Goby, CRAM, SAM or BAM formats. A parameter is used to specify the path to a model to call genotypes.

#### Training new models

Models for new assays can be trained by constructing a training, validation and test datasets. We provide scripts to automate this activity. Detailed steps are documented with the software (see https://github.com/CampagneLaboratory/variationanalysis), but briefly, the datasets are produced by converting alignments to .sbi files with the Goby `SEQUENCE_ BASE_INFORMATION` output format. True genotypes can be introduced in the .sbi file at this step. The .sbi file is randomized and randomly split into training, validation and test sets. The training of the model uses the training and validation sets for early stopping. Final model performance is estimated on the test set to verify that the model generalizes. Models are saved to disk during training.

## Neural Network Architecture

### Feature Mappers

Feature mappers convert alignments about one sample into a fixed set of features suitable for training with neural networks. Regardless of the number of reads aligned at a genomic position, mappers need to produce a fixed-length output so that these outputs can be concatenated consistently into a fixed-length input vector. At each genomic site, a mapper generates the number of reads supporting each genotype (counts), the number of distinct locations in the read that support the genotype (distinct read indices). Hundreds of features are derived for each site and a complete list is provided in the source code. Mappers are implemented in the variationanalysis project available at GitHub https://github.com/CampagneLaboratory/variationanalysis. This study used org.campagnelab.dl.genotype.mappers.GenotypeMapperV13.

### Model Architecture

Models were developed with the DeepLearning4J (DL4J) framework (http://deeplearning4j.org/), version 0.7.1. DL4J was selected because it is a Java framework and the models it produces can be integrated with the Goby framework more easily than frameworks in other languages. Models were formulated as 4 fully connected layers with RELU activation and a fully connected output layers with soft-max activation. The number of output layers depends on the specific label mapper used. The dense inner layers contain 5 times the number of input features. The exact model architecture used is encoded in the class called org.campagnelab.dl.genotype.learning.architecture.graphs.`CombinedWithIsVariantGenotypeAssembler` distributed in the variationanalysis project Torracinta and Campagne [2016].

### Label Mapper

We have experimented with different methods to map genotypes to label vectors. One method calls alleles individually, and encodes the number of alleles. (Implemented with org.campagnelab.dl.genotype.mappers.NumDistinctAllelesLabelMapper and 10 org.campagnelab.dl.genotype.mappers.GenotypeLabelsMapper). This method is suitable for genomes of arbitrary ploidy (e.g., plants). Another method is similar to that described in Poplin et al. [2016] and is limited to diploid genomes.

### Early Stopping

We trained models with early stopping. Briefly, performance of the model was measured on a validation set and training was stopped when performance on the validation set did not increase for 10 epochs. We used the harmonic mean of F1 and AUC as validation performance measure.

### Analysis Protocol Summary

A summary of a typical analysis protocol is provided here. Runnings these steps requires defining some environment variables and is fully explained in the software documentation. Training a model is performed in two high-level steps:

- Transform an alignment (BAM,CRAM or Goby format) into an .sbi file (input for feature and label mappers): `parallel-genotype-sbi.sh 10g NA12878_S1.bam` This step produces the files NA12878_S1-train.sbi and NA12878_S1-train.sbip and two pairs of files, one for validation (used for early stopping) and for test set.
- Train and evaluate the model with a choice of feature mapper (the number 1 is the index of the GPU to train the model on): `iterate-genotype.sh org.campagnelab.dl.genotype.\ mappers.GenotypeMapperV13 1` This produces a model trained with the mapper on the training set and performance statistics on the validation set (printed during training, also stored in the model directory for reference), and finally runs the model on the test set, printing and storing statistics.

## DISCUSSION

We presented a novel approach to call genotypes in high-throughput sequencing data using neural networks. The approach used in this study relies on deep neural networks to call genotypes and can be trained from gold-standard data. It differs from previous approaches in the following ways.

Training deep learning models to call genotypes is a straightforward adaption of some of the ideas that we presented in Torracinta et al. [2016], Campagne [2016]. Given the existence of gold standard data for genotype calls and the ability of deep neural networks to reliably estimate probabilities, the development of a deep learning caller is a logical step. Our method differs from published genotype callers (e.g., GATK) which rely on carefully designed probabilistic models McKenna et al. [2010], Nielsen et al. [2011]. Our approach is to train probabilistic models from data. The key advantage is that new models can be trained and adapted quickly to new experimental assays or data analysis protocols (e.g., combination of aligners and read preprocessing techniques).

Poplin and colleagues have recently demonstrated that a similar idea performs extremely well across a variety of datasets (Poplin et al. [2016]). In this work, these authors have leveraged the expertise of Google in deep learning for images. They converted alignment data to images and trained models with the Inception v2 architecture. Their work is a clear demonstration that deep learning can be used for genotype calling and can result in state of the art genotype calling performance (as measured by genotype concordance, and precision/recall or F1 for the identification of variants).

The approach of Poplin et al has several drawbacks. First, as presented, it only supports calling genotypes for diploid organisms (the network predicts three states for the genotype: AA A/B or BB). A universal caller should also be applicable to plants and other non diploid organisms. Our approach supports and already implements label mappers that can be used with arbitrary ploidy. The computational procedure described in Poplin et al. [2016] has another important drawback: its computational efficiency. Converting alignments to images requires assembling a 10,000 by 300 pixel image for each site where a prediction is required. The DeepVariant model therefore uses at least 3 million pixels per site. In contrast, our approach uses less than a 1,000 floats to represent both features and labels. Since our approach uses orders of magnitude fewer features, the training and evaluation datasets can be stored in a few tens of gigabytes and model training can be conducted on a workstation with one or a few GPU cards (we trained and evaluated models for this study on a workstation costing less than $10K). Computational efficiency is important to allow individual researchers to replicate results, make and evaluate method improvements, and develop models for new experimental assays or analysis protocols.

Our methods also differ from Poplin et al. [2016] in the amount of preprocessing applied to alignment data before it is provided to the neural network. We applied limited preprocessing (realignment around indels), when Poplin et al used the GATK haplotype caller, which implements local reassembly. We believe that many of the pre-processing/clean-up operations currently implemented by ad-hoc software can be trained by back propagation given suitable training data. The fact that the test performance of the models we trained with minimal preprocessing are closing in on state of the art performance suggests that this hypothesis has merit.

Another point of difference between our study and the work of Poplin et al. [2016] is that we distribute the software we developed under an open-source license. The immediate availability of the software and detailed model training protocols will make it possible for other researchers to train models for new platforms as well as to contribute to method development and evaluation.

Finally, we are looking for collaborators interested in helping develop and evaluate improved versions of this genotype caller on a wide range of platforms. Training new models will be required when data from new platforms becomes available and for this reason we feel that a community effort is best suited to efficiently developing these technologies.

The software that we presented here (see also Torracinta et al. [2016], Campagne [2016]) provides a test-bed infrastructure where new ideas can be tested and evaluated quickly. We hope that it will enable a community of researchers to experiment with neural networks for genomic applications.

## ACKNOWLEDGMENTS

We thank Manuele Simi for technical assistance with Maven configurations for this project.

## REFERENCES

Fabien Campagne. http://dx.doi.org/10.1101/079087 continuation: Evaluation of adaptive somatic models in a gold standard whole genome somatic dataset. bioRxiv, 2016. doi: 10.1101/093534. URL http://biorxiv.org/content/early/2016/12/13/093534.

Fabien Campagne, Kevin C. Dorff, Nyasha Chambwe, James T. Robinson, and Jill P. Mesirov. Compression of Structured High-Throughput Sequencing Data. PLoS ONE, 8(11):e79871, nov 2013. ISSN 1932-6203. doi: 10.1371/journal.pone.0079871. URL http://dx.plos.org/10.1371/journal.pone.0079871.

Michael A Eberle, Epameinondas Fritzilas, Peter Krusche, Morten Källberg, Benjamin L Moore, Mitchell A Bekritsky, Zamin Iqbal, Han-Yu Chuang, Sean J Humphray, Aaron L Halpern, et al. A reference data set of 5.4 million phased human variants validated by genetic inheritance from sequencing a three-generation 17-member pedigree. Genome Research, 2016.

Markus Hsi-Yang Fritz, Rasko Leinonen, Guy Cochrane, and Ewan Birney. Efficient storage of high throughput dna sequencing data using reference-based compression. Genome research, 21(5):734–740, 2011.

Heng Li, Bob Handsaker, Alec Wysoker, Tim Fennell, Jue Ruan, Nils Homer, Gabor Marth, Goncalo Abecasis, Richard Durbin, et al. The sequence alignment/map format and samtools. Bioinformatics, 25 (16):2078–2079, 2009.

Aaron McKenna, Matthew Hanna, Eric Banks, Andrey Sivachenko, Kristian Cibulskis, Andrew Kernytsky, Kiran Garimella, David Altshuler, Stacey Gabriel, Mark Daly, et al. The genome analysis toolkit: a mapreduce framework for analyzing next-generation dna sequencing data. Genome research, 20(9): 1297–1303, 2010.

Rasmus Nielsen, Joshua S Paul, Anders Albrechtsen, and Yun S Song. Genotype and snp calling from next-generation sequencing data. Nature Reviews Genetics, 12(6):443–451, 2011.

Ryan Poplin, Dan Newburger, Jojo Dijamco, Nam Nguyen, Dion Loy, Sam S. Gross, Cory Y. McLean, and Mark A. DePristo. Creating a universal snp and small indel variant caller with deep neural networks. bioRxiv, 2016. doi: 10.1101/092890. URL http://biorxiv.org/content/early/2016/12/21/092890.

Remi Torracinta and Fabien Campagne. Variationanalysis 1.0.2 software release, October 2016. URL https://doi.org/10.5281/zenodo.159203.

Remi Torracinta, Laurent Mesnard, Susan Levine, Rita Shaknovich, Maureen Hanson, and Fabien Campagne. Adaptive somatic mutations calls with deep learning and semi-simulated data. bioRxiv, 2016. doi: 10.1101/079087. URL http://biorxiv.org/content/early/2016/10/04/079087.

